# EXONIZATION BY THE EMERGENCE OF A CLEAVAGE-POLYADENYLATION SITE

**DOI:** 10.1101/2022.03.17.484714

**Authors:** Agnès Méreau, Hubert Lerivray, Justine Viet, Serge Hardy, Luc Paillard, Yann Audic

**Affiliations:** Univ Rennes, CNRS, IGDR (Institut de Génétique et Développement de Rennes), UMR 6290, F-35000 Rennes, France

**Keywords:** exonization, 3’ terminal exons, alternative splicing, amphibians, Xenopus

## Abstract

Exonization is the evolutionary process of recruitment of new exonic regions from previously intronic regions. It is a major contributor to the increased complexity of alternative splicing. Here, we explore exonization mediated by the emergence of a novel cleavage-polyadenylation site in an intron. In *Xenopus laevis*, the *tpm1* gene, which encodes muscular tropomyosin, contains alternative terminal exons. In adult muscles and embryonic hearts, exon 9A is joined to the terminal exon 9B. In embryonic somites, it is joined to the exonic region 9’, which is transcribed from the intron immediately downstream of exon 9A. Consequently, exon 9A is either an internal exon when ligated to exon 9B, or a part of a terminal exon along with region 9’. We show here that region 9’ is present only in amphibians and coelacanths. This suggests that it emerged in sarcopterygians and was lost in amniotes. We used antisense morpholino oligonucleotides to mask the regions of *tpm1* pre-mRNA that potentially regulate the inclusion of exon 9A9’. This revealed that the definition of exon 9A9’ relies on a weak cleavage-polyadenylation site and an intronic enhancer, but is independent of the 3’ splice site. We demonstrate that RNAs containing exon 9B are toxic in somites. This may have contributed to the evolutionary pressure that led to the exonization of region 9’ in sarcopterygians. These findings reveal the emergence of a novel cleavage-polyadenylation site that avoids the accumulation of a toxic RNA as a novel mechanism for exonization-mediated diversification of terminal exons.

## INTRODUCTION

Splicing is the process of removing introns and ligating exons of pre-RNAs to produce mature RNA molecules, most often mRNAs encoding proteins. The key signals for splicing are the splice donor or 5’ splice site, the splice acceptor or 3’ splice site, and the branch point located within the intron generally close to the 3’ splice site. Terminal exons are specifically delineated at their 3’ end by a cleavage and polyadenylation site. During the first step of splicing, the 5’ splice site is ligated to the branch point. During the second step, the released 3’ end of the upstream exon is ligated to the downstream 3’ splice site (5’ of the downstream exon). This results in creation of an exon junction and release of the intron (Wan et al. 2020; Wilkinson et al. 2020). During alternative splicing, the conditional inclusion or skipping of a subset of exons allows for the production of several different mRNA molecules from a single gene, potentially encoding different proteins. A systematic comparison of the genomes and transcriptomes of 47 eukaryotic species revealed a strong relationship between organism complexity (number of cell types) and alternative splicing, whereas the number of protein-coding genes in an organism only poorly correlated with its complexity (Chen et al. 2014). For example, in the budding yeast *Saccharomyces cerevisiae*, deep sequencing approaches have revealed that only 306 protein-coding genes out of 6000 contain introns, with alternative 5’ or 3’ splice sites in 20% of them (Schreiber et al. 2015). Conversely, more than 92% of human genes produce at least two alternative mRNA isoforms (Pan et al. 2008; Wang et al. 2008). Interestingly, within vertebrates, the splicing profiles of the same organs strongly diverge between species (Barbosa-Morais et al. 2012). During vertebrate evolution, the tissue-specificity of gene expression has been largely conserved, whereas the alternative splicing patterns across species have been conserved only for a subset of tissues (Merkin et al. 2012).

A major force driving the increasingly complex nature of alternative splicing patterns during evolution is the appearance of novel exons. They can originate from the “exonization” of previously intronic regions (Parma et al. 1987; Keegan et al. 2022). In this process, mutations lead to the appearance of splice sites within introns. The corresponding intronic regions are then retrieved as exons within the mature mRNAs. The mutations that lead to exonization are generally well tolerated because the new splice sites are usually less efficient than the canonical ones, and the new exon is only incorporated into a small proportion of the mature mRNA molecules. Over the course of evolution, selection of further mutations may strengthen the new splice sites if the incorporation of the new exon is advantageous (Schmitz and Brosius 2011).

Transposable elements are a considerable source of intronic regions with the capacity to undergo exonization. Systematic comparison of transposable elements in chick, zebrafish, and non-vertebrate species revealed more exonization in vertebrates. All types of transposable element, including long interspersed nuclear elements (LINE) and short interspersed nuclear elements (SINE), are susceptible to exonization (Krull et al. 2005; Sela et al. 2010). Alu elements are 300 nucleotide-long primate-specific SINEs that make up 10% of the human genome. Alu elements greatly contributed to expanding alternative splicing patterns in primates by providing novel exons. They can also cause diseases such as neurofibromatosis type 1 where an Alu insertion modifies the splicing of *NF1* pre-mRNA (Wallace et al. 1991). Generally, Alu-containing exons are alternative rather than constitutive exons (Sorek et al. 2002). Alu-containing exons are enriched in 5’ untranslated regions, where their alternative inclusion or skipping modulates mRNA translation (Shen et al. 2011). They are also found immediately downstream of genes, promoting alternative terminal exons. This results in proteins with different C-terminal domains and/or mRNAs with different 3’ untranslated regions (3’UTR), with a potential impact on mRNA translation or stability (Tajnik et al. 2015). An Alu-containing exon in the glycoprotein hormone alpha (GPHA) gene modifies the N-terminal region of the encoded protein. This increases the serum half-life of human chorionic gonadotropin, of which GHPA is a subunit (Chen et al. 2017). A key event for Alu exonization is the appearance of a branch point upstream of the novel exon, which is recognized by the U2AF65 protein. The RNA-binding protein hnRNPC competes with U2AF65 for potential new branch points, and therefore limits the consequences of Alu exonization (Zarnack et al. 2013; Tajnik et al. 2015).

Exonization of a transposable element that lies within an intron provides new cassette exons, but this does not explain other types of alternative splicing. Notably, the emergence of alternative 5’ or 3’ splice sites constitutes a mechanism to exonize an intronic region immediately adjacent to an existing exon (Koren et al. 2007). Another potential mechanism would be the emergence of a cleavage and polyadenylation site in the intron downstream of an internal exon. However, to our knowledge, this has never been investigated in detail. The *tpm1* gene in *Xenopus laevis* encodes tropomyosin alpha-1, a major component of the cytoskeleton. Muscular and non-muscular isoforms of *tpm1* mRNA differ in their terminal exons. In non-muscular isoforms, exon 8 is ligated to a distant terminal exon, 9D. In muscular isoforms, exon 8 is ligated to exon 9A (Fig. 1A). In embryonic hearts and in adult muscles, exon 9A is ligated to terminal exon 9B. While muscular and non-muscular isoforms of *TPM1* are found in all vertebrates, a third isoform exists in *X. laevis*. In this species, the intron downstream of exon 9A contains an alternative cleavage-polyadenylation site, which is used in embryonic somites (presumptive skeletal muscles). The region between the 5’ splice site and the cleavage-polyadenylation site is named 9’. Consequently, in addition to a non-muscular isoform (8-9D), two alternative muscular isoforms exist in *X. laevis* (Fig. 1A), isoform 8-9A-9B and isoform 8-9A9’ (Hamon et al. 2004). Here, we take exon 9’ as an example of an intronic region that has been exonized by the emergence of a cleavage and polyadenylation site within a previously intronic region. We show that exon 9’ appeared in sarcopterygians after their divergence from ray-finned fishes, and was lost in amniotes. We decipher the complex regulation leading to the inclusion of exon 9’ in mature mRNAs, and we demonstrate that its usage relies on an AG-independent 3’ splice site and on an activating cis-element in the upstream intron. Finally, we tackle why the emergence of exon 9’ was advantageous and evolutionarily selected by demonstrating that the ectopic expression of exon 9B is toxic for *Xenopus* embryos. Together, our results reveal a new mechanism that leads to the appearance of a novel exonic region during evolution, contributing to alternative splicing pattern diversity in vertebrates.

**Figure 1.**
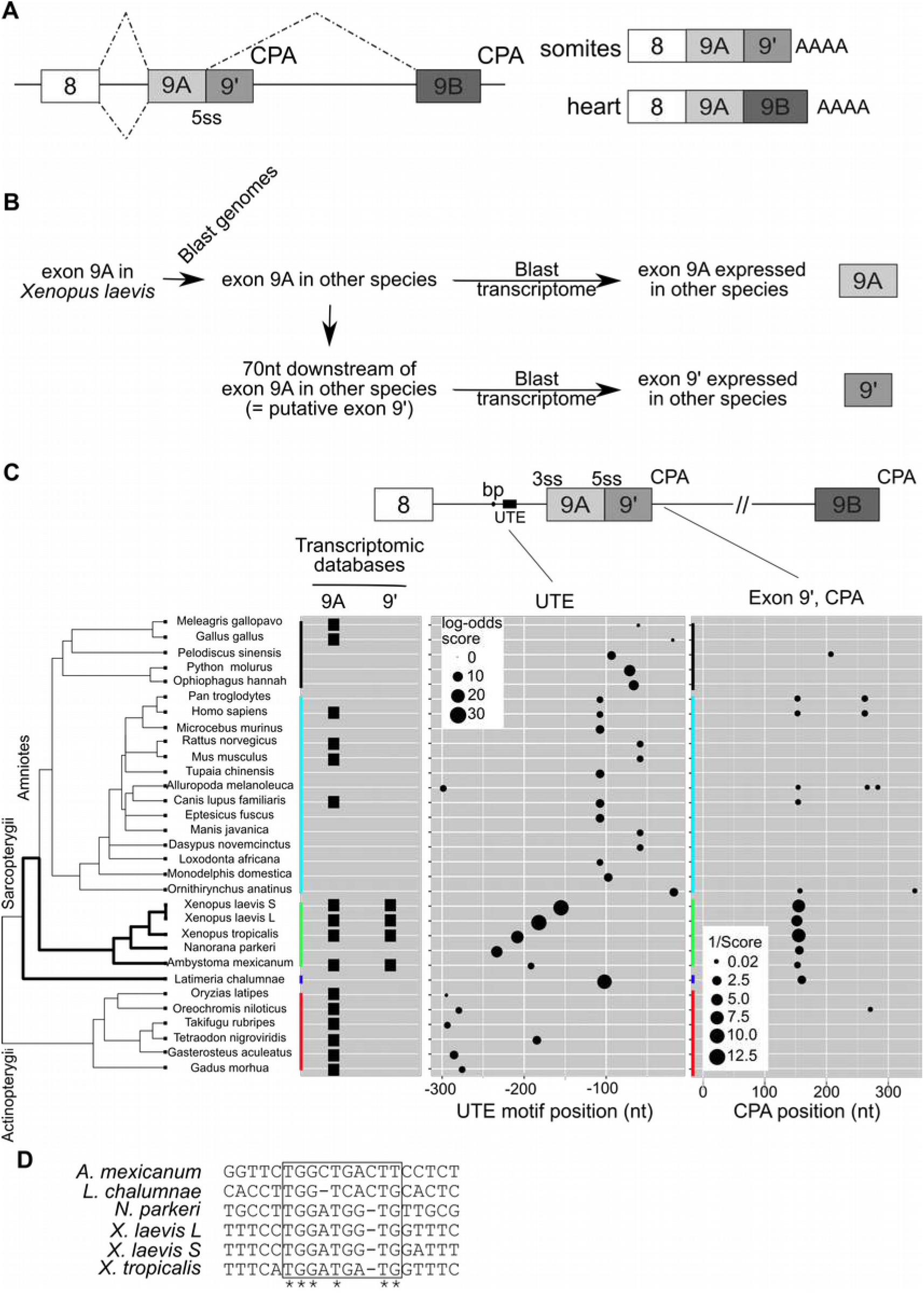
Genomic region and regulatory elements delineating *TPM1* terminal exons. **A**, Schematic of Xenopus muscular *tpm1* terminal exons and alternative, tissue-specific, maturation patterns. In embryonic heart, exon 9A is joined to exon 9B. In somites, the 5’ splice site of the intron between exons 9A and 9B (5ss) is not used, and cleavage and polyadenylation (CPA) occur 75 nucleotides downstream of exon 9A. The region between the 5ss and the CPA site is referred to as 9’. **B**, Strategy to identify *TPM1* genes and the species that express the genomic region 9’. **C**, Left, phylogenetic tree from NCBI taxonomy database showing the relationships between the species that were analyzed. Right, the results of transcriptomic database (EST) searches for regions 9A and 9’, the results of UTE sequence scoring of the 300 nucleotides upstream of exon 9A and the results of CPA scoring of the 300 nucleotides downstream of exon 9A. The size of the circles is proportional to the predicted strength of the site. **D**, Alignment of UTE sequences in amphibians and coelacanth (Clustal Omega).

## RESULTS AND DISCUSSION

### Emergence and loss of the *TPM1* exon 9A9’

Two alternative muscular RNA isoforms of *tpm1* exist in *X. laevis* heart and somites. They differ by the use of either a 5’ splice site (5ss) or of a cleavage and polyadenylation site (CPA) downstream of exon 9A (Fig. 1A). Except for the final amino acid, the encoded tpm1 proteins are the same, and the two isoforms differ only in their 3’UTR (Hardy et al. 1999). We wondered when and how exon 9’ emerged, as it has not been described in any species other than *X. laevis*. We used the strategy shown in Fig. 1B to investigate whether a similar exonic organization exists in other vertebrate species. Using Blastn and Genomicus (Nguyen et al. 2018), we identified the *TPM1* orthologs in 30 vertebrate species based on synteny. We collected *TPM1* genomic sequences and annotated exons 8, 9A and 9B according to peptide or nucleotide homology using tblastn and cDNA alignment. We checked whether genomic region 9A is transcribed by looking for its sequence in transcriptomic databases. We retrieved the 70-nucleotide genomic sequence downstream of exon 9A, corresponding to the putative exonic region 9’, and searched for it in transcriptomic databases.

Genomic regions 9A and 9’ were found in the transcriptomes of amphibians such as frogs (*X. laevis, X. tropicalis*) and salamanders (*Ambystoma mexicanum*). Exon 9A, but not putative exon 9’, is found in transcriptomic databases of several ray-finned fishes (actinopterygians) and amniotes (Fig. 1C, left panel). This strongly suggests that no CPA immediately downstream of exon 9A is used, and that only the isoform containing exons 9A and 9B is present in muscles in these species. This analysis suggests that exon 9’ is amphibian-specific.

To understand why exon 9A9’ is apparently used as a terminal exon only in amphibians, we examined all the genomic cis-sequences that are potentially involved in its regulation. We previously found that, in *X. laevis*, an intronic enhancer, UTE, is located upstream of exon 9A and targets the cleavage/polyadenylation machinery to the CPA in exon 9’ (Anquetil et al. 2009). We combined two approaches to investigate whether this element is present upstream of exon 9A in other species. Firstly, we scored the UTE motif defined from the two *X. laevis* UTE sequences

(corresponding to the two *tpm1* homoeologous genes) in the 300 intronic nucleotides upstream of exon 9A (Fig. 1C, middle panel). High UTE scores were obtained for *Xenopus laevis* and *Xenopus tropicalis, Nanorana parkeri, Latimeria chalumnae* (coelacanth), *Python molurus* and *Ophiophagus hannah* (two snakes). Multiple sequence alignment (Clustal Omega) identified a further sequence element in *A. mexicanum* that shares a high conservation with the *Xenopus* core UTE (Fig. 1D).

We next used CPA scoring to investigate whether a cleavage and polyadenylation site is located downstream of exon 9A (Cheng et al. 2006). Strong CPA sites were found in amphibians, but also in *L. chalumnae* and to a lesser extent in some mammals including human, but not in snakes (Fig. 1C, right panel). For other sequence elements likely to control exon 9A9’ usage, we found no difference between amphibians and other species. This includes the strength of the branch point within intron 8, the 5’ splice site between exons 9A and 9’, the branch point within intron 9 (upstream of exon 9B) or the cleavage-polyadenylation site of exon 9B (Fig. S1). Hence, the utilization of exon 9A9’ as a terminal exon is associated with the simultaneous occurrence of the UTE and the cleavage-polyadenylation site in exon 9’.

We could not assess the existence of a 9’ sequence in mRNAs from *L. chalumnae* (coelacanth) or *Nanorana parkeri* (Tibetan frog) due to the lack of published transcriptomic data. However, these two species have both a UTE and a cleavage-polyadenylation site (Fig. 1C), strongly suggesting that the 9’ region can be included in their *tpm1* mRNA. Therefore, we experimentally tested whether genomic sequences surrounding exon 9A from *L. chalumnae* and *N. parkeri* can support the use of exon 9A9’ as a terminal exon. We injected *Xenopus* embryos with a plasmid (Fig. 2A) containing the *X. laevis* genomic region between exons 7 and 9B under the transcriptional control of a somite-specific promoter (Hamon et al. 2004). We replaced the *Xenopus* region 9A9’ by the equivalent regions from *L. chalumnae* or *N. parkeri*, or from *G. gallus* as a negative control. We extracted RNAs from tailbud embryos, and analyzed the splicing pattern of the mRNA transcribed from the injected plasmid by RT-PCR. One PCR primer is specific to the plasmid sequence and cannot amplify endogenous *tpm1* mRNA (Fig. 2B, lanes 1 and 8). As previously described (Hamon et al. 2004), three mRNA isoforms are produced from the injected *Xenopus* reporter gene (Fig. 2B, lanes 2 and 9). As expected, inactivating the 3’ splice site by mutation of the AG to GA (d3’), or strengthening the weak internal 5’ splice site by creating a consensus site leads to the exclusion of exon 9A or the strict inclusion of exon 9A as an internal exon, respectively (lanes 3, 4, 10, 11). When we replaced the *X. laevis* genomic region surrounding exons 9A9’ in the splicing reporter gene by its *N. parkeri* equivalent, the compound minigene produced the same mRNA isoforms as the original *X. laevis* gene (Fig. 2B, lanes 5 and 12). Notably, a 7-8-9A9’ isoform was detected (lane 12), demonstrating that the genomic region around exon 9A in *N. parkeri* efficiently drives the utilization of terminal exon 9A9’. Two other isoforms were also produced (highlighted by asterisks in Fig. 2B). They originate from a cryptic 5’ splice site in *N. parkeri* intron 8 (Fig. S2A-B). Strikingly, the *L. chalumnae* compound reporter minigene produced mainly the 7-8-9A9’ mRNA isoform (Fig.2B, lanes 6 and 13). Consistent with the lack of a 9’ terminal exon, the lack of UTE and the lack of a potential CPA in the chicken genome (Fig. 1C), the *X. laevis* - *G. gallus* compound minigene produced no mRNA isoform containing the 9’ region (Fig. 2B, lane 14). All the splicing reporter minigenes were transcribed with similar efficiencies, as revealed by RT-PCR targeting constitutive exons 7 and 8 (Fig. S2C). Together, these data show that *N. parkeri* and *L. chalumnae* genomic sequences can drive the maturation of *tpm1* pre-mRNA toward isoforms containing the exonic region 9’ as a terminal exon. The most parsimonious interpretation of an exon 9A9’ in amphibians and coelacanths but not in other species is that this terminal exon emerged in sarcopterygians following the divergence from actinopterygians (ray-finned fishes), and was lost in amniotes (Fig. 1C, left).

**Figure 2:**
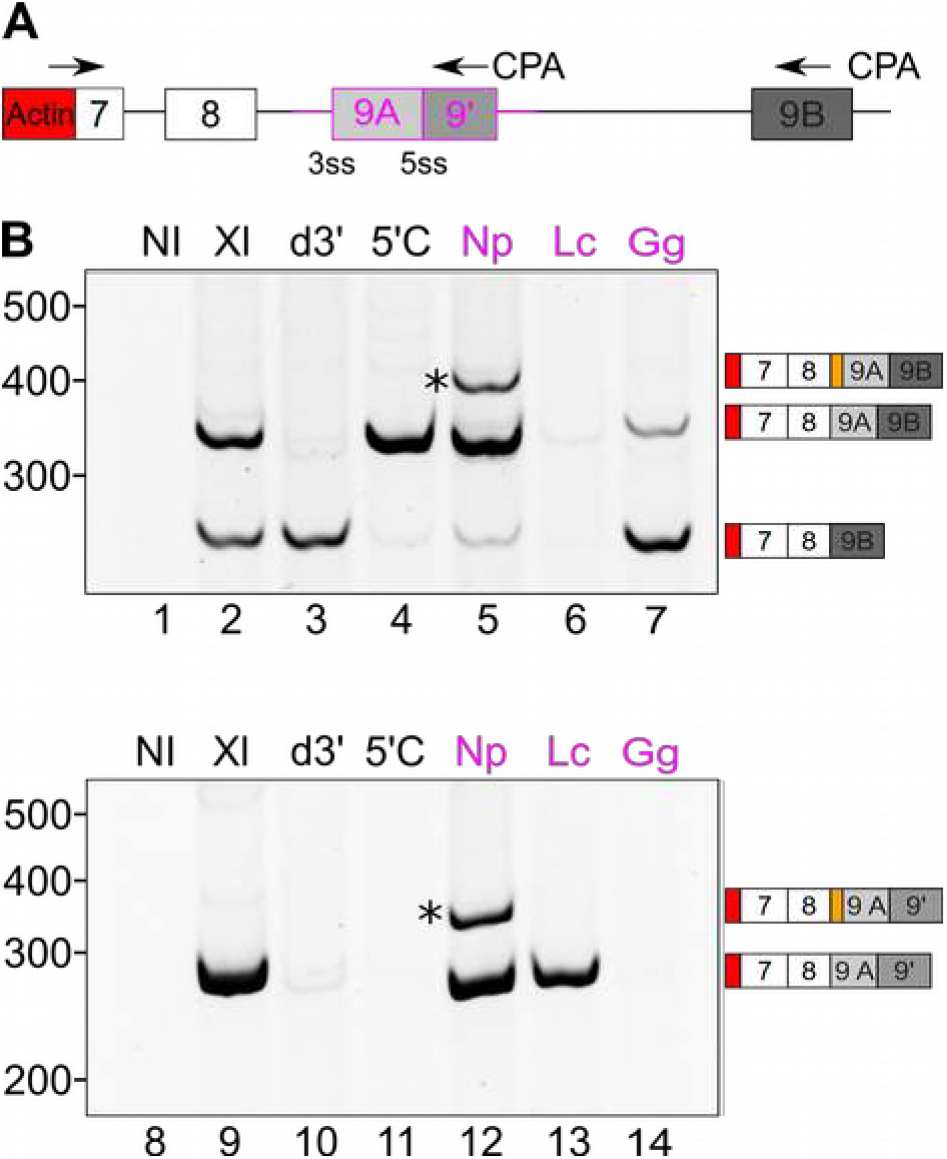
The amphibian and coelacanth 9A9’ is used as a terminal exon. **A**, Splicing reporter assay. The splicing reporter gene consists of the *X. laevis* genomic region between exons 7 and 9B cloned downstream of the actin promoter. In compound genes, the original *X. laevis* region covering exon 9A9’ and a part of the flanking introns (purple box and lines, approximately 1000 nucleotides) is replaced by the orthologous region from other species (Np, *Nanorana parkeri*, Lc *Latimeria chalumnae*, Gg *Gallus gallus*). RT-PCR primers used for the analysis of the splicing patterns prime at the promoter-exon 7 junction, in exon 9’ and in exon 9B (arrows). **B**, Following injection into embryos, the maturation patterns of the RNAs produced from this reporter gene were analyzed by RT-PCR with the forward and reverse primer targeting exon 9B (upper panel) or exon 9’ (lower panel). Lanes 1 and 8, non-injected embryos. Lanes 2 and 9, *Xenopus* reporter gene. Lanes 3 and 10, *Xenopus* gene with inactive intron 8 3’ splice site (d3’). Lanes 4 and 11, *Xenopus* gene with strengthened 5’ splice site at the 9A-9’ junction (5’C). Lanes 5-7 and 12-14, compound genes. The identities of all bands were confirmed by sequencing.

### The UTE functionally replaces the 3’ splice site to define terminal exon 9A9’

The above data reveal that in sarcopterygians, an internal exon gained the ability to be used as a terminal exon. This new property relies on the appearance of a CPA downstream of exon 9A and of a UTE upstream of exon 9A. The elucidation of the underlying molecular mechanisms is a key step to understand how this splicing event has been phylogenetically conserved. We used *X. laevis* to delve deeper into the regulation of 9A9’ as a terminal exon.

Exon definition is prevalent in vertebrates. In this model, the future exon is defined by the formation of a cross-exon complex linked to the two ends of the exon (Guth et al. 1999; Wu et al. 1999; Guth et al. 2001). To check whether exon 9A9’ is accurately described by this model *in vivo*, we used antisense morpholino oligonucleotides to block either of the two ends of exon 9A9’ of endogenous *tpm1* pre-mRNA. In non-injected or control morpholino-injected embryos, three isoforms were detected (Fig. 3A, lanes 1-2). As expected, blocking the 3’ end of exon 9A9’ with a morpholino that targets the cleavage and polyadenylation site (pAMO) reduced the abundance of the isoform that uses this CPA (8-9A9’) in favor of an isoform that does not use it (8-9A-9B, lane 3). Blocking the 5’ end of exon 9A9’ with a morpholino that targets the 3’ splice site (3ssMO) reduced the abundances of the two isoforms that use this 3’ splice site (8-9A9’ and 8-9A-9B, lane 5). However, the reduced abundance of these two isoforms was not accompanied by an increase of isoform 8-9B. This apparently contradicts the exon definition model, which states that impairing the definition of an exon results in exon skipping (Guth et al. 1999; Wu et al. 1999; Guth et al. 2001).

**Figure 3.**
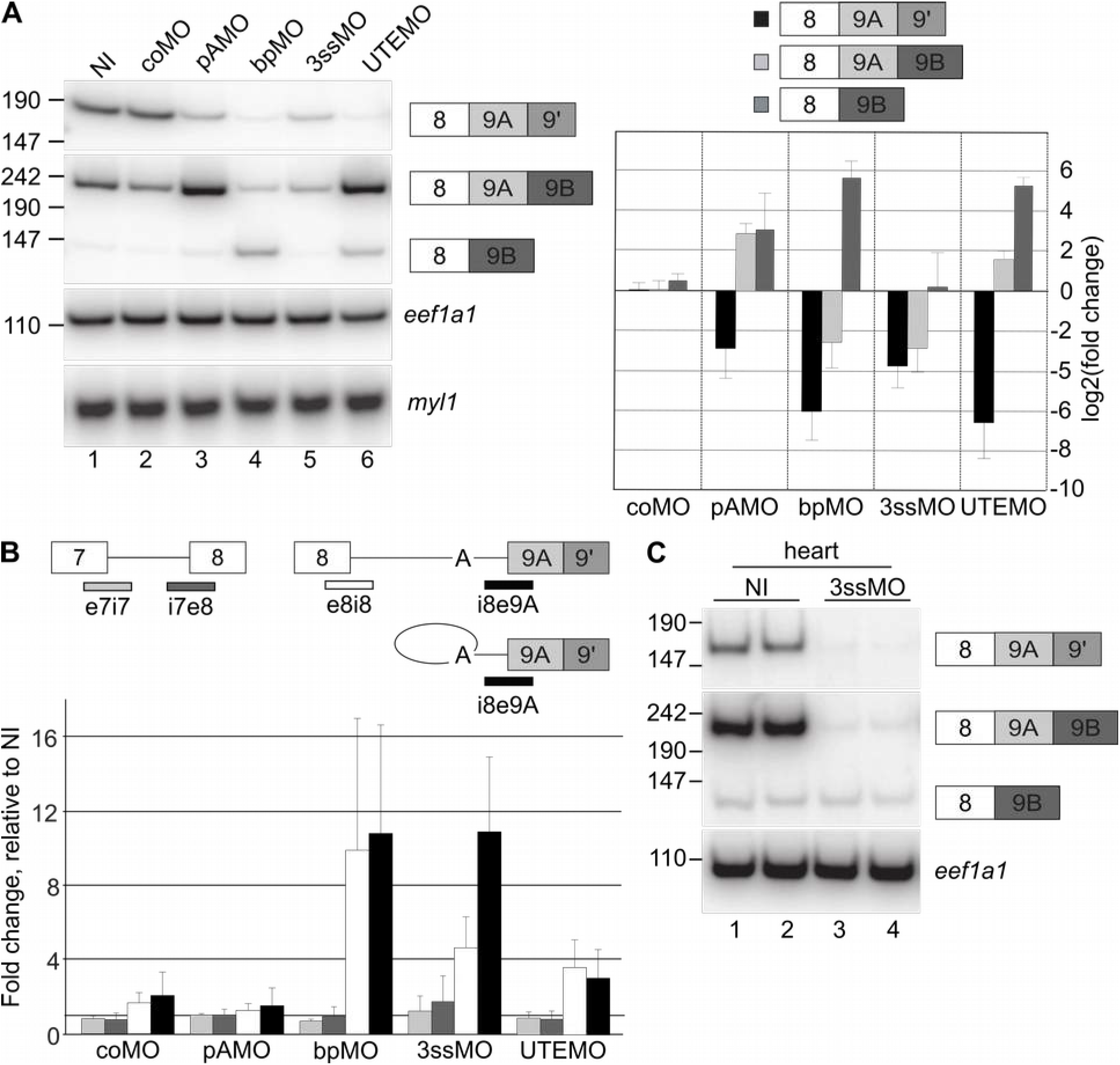
The definition of terminal exon 9A9’ is independent of the 3’ splice site but requires the UTE. We injected embryos with morpholinos and allowed them to develop until stage 28. coMO, control morpholino; pAMO, morpholino against the CPA in exon 9’; bpMO, morpholino against the branch point in intron 8; 3ssMO, morpholino against the 3’ splice site of intron 8; UTEMO, morpholino against the UTE; NI, not injected. **A**, Left panel, RT-PCR using a forward primer in exon 8 and reverse primers in exon 9’ (upper panel) or 9B (middle panel). RNA recovery and embryo stage were assessed by amplification of *eef1a1* and *myl1* mRNA respectively. Right panel, quantification of at least 3 independent experiments, relative to the non-injected embryos (mean ± standard deviation). **B**, Real-time RT-PCR with primers in exon 7 and intron 7 (light grey, e7i7), in intron 7 and exon 8 (dark grey, i7e8), in exon 8 and intron 8 (white, e8i8) or in intron 8 and exon 9A (black, i8e9A). Fold changes are relative to the non-injected embryos and are presented as mean ± standard deviation of at least 3 independent experiments. **C**, as **A**, using RNA from dissected hearts.

In the absence of an in-frame stop codon, the endogenous isoform 8-9B is subject to non-stop decay (Anquetil et al. 2009). Its instability could explain why we failed to detect an increase of this isoform following the injection of the 3ssMO. Therefore, we decided to block the branch point within intron 8 as an alternative approach to inhibit the definition of exon 9A9’. This required the identification of the branch point *in vivo*. To do so, we used an RT-PCR based method (Vogel et al. 1997) utilizing a back-to-back pair of primers that amplify a fragment of intron 8 across the branch point in the lariat (Fig. S3A). The size and the sequence of the amplimere indicate that the branch point is at the adenine nucleotide located 274 nucleotides upstream of the 3’ splice site (Figs. S3B-C). This is consistent with *in silico* predictions (Fig. S1). Although this branch point is quite distant (branch points are generally 20 to 50 nucleotides upstream of the 3’ splice site), this is in line with the presence of a long (21 nucleotides) pyrimidine tract immediately downstream, and the absence of AG dinucleotide between it and the 3’ splice site (Fig. S1). We used this information to design a morpholino antisense oligonucleotide that blocks the branch point (bpMO). Upon injection, this morpholino strongly reduced the abundance of both isoforms 8-9A9’ and 8-9A-9B (Fig. 3A, lane 4), demonstrating that it efficiently impedes the definition of exons 9A and 9A9’. Importantly, it strongly increased the abundance of the isoform 8-9B. This demonstrates that an increased abundance of isoform 8-9B can be observed despite its instability. In contrast, the fact that the isoform 8-9B was not increased by the morpholino against the 3’ splice site (lane 5) indicates that the 3’ splice site is not required for exon 9A9’ definition, even if an accessible 3’ site is required for the ligation of exon 8 to exon 9A9’.

If exon 9A9’ is defined independently of the 3’ splice, then preventing the recognition of the 3’ splice site should only inhibit the second trans-esterification reaction, leading to the accumulation of introns with non-excised lariats. To test this hypothesis, we developed a RT-qPCR assay to quantify exon-intron boundaries (Fig. 3B). As a control, we checked that neither the exon 7/intron 7 (e7i7) nor the intron 7/exon 8 (i7e8) boundaries were affected by the injected molecules (Fig. 3B, light and dark grey, respectively). In contrast, the bpMO strongly increased the abundance of both exon 8/intron 8 (e8i8) and intron 8/exon 9A (i8e9A) boundaries (Fig. 3B, white and black, respectively). This confirms that blocking the recognition of the intron 8 branch point inhibits the excision of this intron. Importantly, the 3ssMO increased the abundance of i8e9A much more efficiently than that of e8i8. Amplification of e8i8 only occurs from *tpm1* pre-mRNA, while amplification of i8e9A also occurs from the RNA species containing non-excised lariat. Hence, the 3ssMO caused the accumulation of non-excised lariat RNA. This supports the interpretation that exon 9A9’ is defined independently of the 3’ splice site. In addition, while the isoform 8-9A9’ is predominant in total embryos, the isoform 8-9A-9B is predominant in dissected hearts (Fig. 3C) (Hardy et al. 1999). In hearts, the 3ssMO reduced the abundance of both 8-9A9’ and 8-9A-9B isoforms (lanes 3-4), but without increasing the abundance of isoform 8-9B. Therefore the internal exon 9A is defined independently of the 3’ splice site, in the same way as the terminal exon 9A9’. Together, these data suggest that intron 8 has a distant branch point with a long pyrimidine tract, and that the definition of exons 9A or 9A9’ does not rely on the recognition of the AG 3’ splice site by U2AF1 (U2AF35). Intron 8 can therefore be described as AG-independent (Guth et al. 1999; Wu et al. 1999; Guth et al. 2001).

The above data raise the question of how the 5’ end of exons 9A and 9A9’ is defined. To test whether the UTE is involved in this process, we injected embryos with a morpholino that masks the UTE (UTEMO) and allowed them to develop as above. The UTEMO dramatically reduced the abundance of isoform 8-9A9’ (Fig. 3A, lane 6). Conversely, it did not reduce the abundance of isoform 8-9A-9B. Together, these data suggest that the UTE is required to define exon 9A9’ as a terminal exon, but not exon 9A as an internal exon.

We used an RNase protection assay to confirm these findings by examining the accumulation of RNA maturation intermediates (Fig. 4). The probe directed against *myl1* mRNA controls RNA recovery and embryo staging. The probe against *tpm1* spans exon 9A9’ as well as a part of the flanking introns. With this probe, the identity of the maturation intermediates can be deduced from the size of the protected fragments (Fig. 4A, upper panel). The probe is directed against the *tpm1*.*L* homoeologous gene, and hybridization with the *tpm1*.*S* RNA resulted in a supplemental cleavage within exon 9’ due to a single nucleotide variation between the 2 homoeologous genes over the probe region. To ensure clarity of the analysis, we only considered the protected fragments arising from the *tpm1*.*L* RNA (colored lines in Fig. 4). One representative experiment is shown in Fig. 4A, replicates are shown in Fig. S4, and the quantification of independent experiments is shown in Fig. 4B. As expected, the protected fragments from non-injected and control MO-injected embryos (lanes 3-4) essentially arose from the hybridization of the probe with the mature RNAs (8-9A9’, 152 and 129 nucleotides, and 8-9A-9B, 79 nucleotides). The pAMO increased the abundance of the 8-9A-9B isoform (79 nucleotides), consistent with previous findings (Fig. 3A), but also of an intermediate matured at its 5’ end but not its 3’ end (216 nucleotides, lane 7). Conversely, in 3ssMO-injected embryos, we detected an intermediate matured at its 3’ end but not its 5’ end (185 nucleotides, lane 6). Importantly, in embryos injected with a morpholino directed either against the branch point or the UTE (lanes 5 and 8), the fully non-processed RNA (249 nucleotides) was predominant due to impaired exon 9A9’ definition. However, while mature isoform 8-9A-9B production (79 nucleotides) was strongly reduced by the morpholino against the branch point, it was not affected by the morpholino against the UTE. This confirms that the UTE is required to define terminal exon 9A9’, but not internal exon 9A. In addition, these results reveal that inhibiting the UTE abrogates both splicing of exon 8 to exon 9A and cleavage at the 3’ end of exon 9’. Hence, exon 9A9’ must be defined at both its 5’ end and its 3’ end before maturation onset, and the UTE is required to couple splicing with cleavage and polyadenylation.

**Figure 4.**
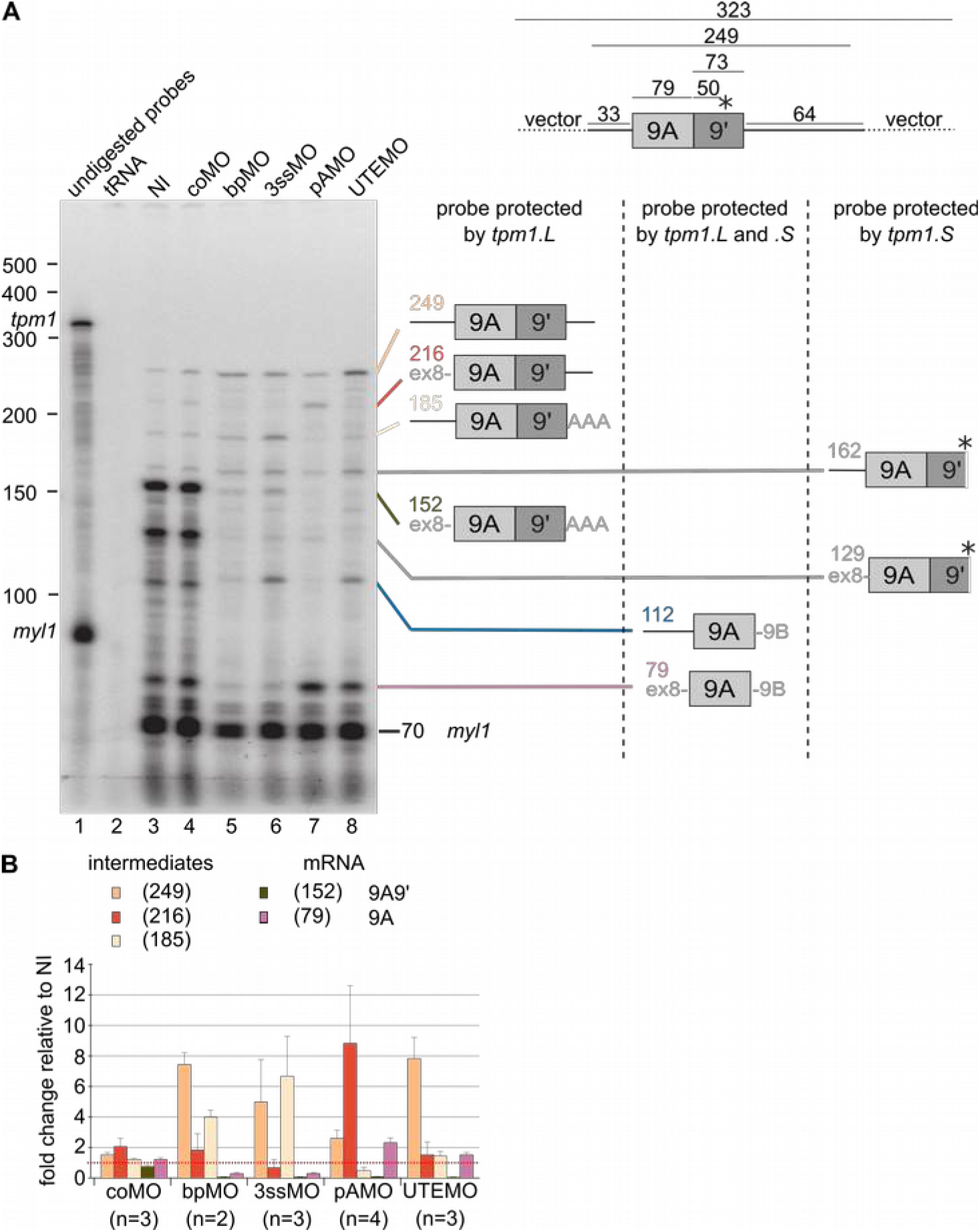
The UTE couples exon 9A9’ splicing and cleavage. **A**, Left, representative RNase protection assay of RNAs extracted from tadpoles previously injected with the indicated morpholinos: coMO, control morpholino; pAMO, morpholino against the CPA in exon 9’; bpMO, morpholino against the branch point in intron 8; 3ssMO, morpholino against the 3’ splice site of intron 8; UTEMO, morpholino against the UTE; NI, not injected. Lane 1, undigested probe mix, with the positions of the undigested probes for *tpm1* and *myl1* indicated on the left. Right, schematic drawing of the 9A9’ region and the probe used for the assay, with the sizes of the protected fragments. The colored lines and numbers indicate the positions of the probe fragments that were protected by *tpm1*.*L* RNA (fully complementary to the probe). The grey lines indicate the positions of probe fragments protected by *tpm1*.*S* RNA (with a mutation in the 9’ region), which were not taken into account for quantification. **B**, Quantification of experiment shown in **A**.

### A driving force for the emergence of *tpm1* exon 9A9’ in sarcopterygians

The above results show that the 3’ splice site of intron 8 is AG-independent, indicating that U2AF is recruited to the strong pyrimidine tract to define exon 9A or 9A9’ independently of U2AF1 binding to the AG dinucleotide. The UTE is required to define exon 9A9’, but not exon 9A. When the UTE is inactive, in heart or following UTEMO injection, 5’ splice site use is predominant. This suggests that the cleavage-polyadenylation site is intrinsically weak. Conversely, in somites, where the UTE is active, it strongly reinforces the efficiency of cleavage-polyadenylation, leading to the production of the 8-9A9’ isoform. Consequently, after their divergence from ray-finned fishes (see Fig. 1C), sarcopterygians evolved a two-pronged mechanism to produce a muscular tropomyosin isoform devoid of the ancestral 3’UTR (exon 9B): an intrinsically weak cleavage-polyadenylation site, and a sequence element within the upstream intron (the UTE), which conditionally defines a terminal exon with an alternative 3’UTR (9’) in somites. This raises the question of the constraints that led to the appearance of this novel 3’UTR in muscular tropomyosin mRNA.

3’UTRs are the site of most translational controls, and we first hypothesized that the two alternative 3’UTRs confer different translational efficiencies on *tpm1* mRNA. We injected *Xenopus* embryos with the pAMO, as it efficiently switches *tpm1* maturation pattern from mainly 8-9A9’ to 8-9A-9B (Fig. 3A lane 3). We assessed the impact of 3’UTR switching on tpm1 protein accumulation by western blot. Injecting the pAMO only weakly reduced muscular tropomyosin accumulation. A much more efficient depletion of tpm1 was achieved by injecting a translation-inhibiting morpholino (Fig. 5A). Furthermore, in luciferase reporter assays, the 3’UTRs 9B and 9’ did not differentially affect the translation of a reporter gene (Fig. S5). We therefore consider it unlikely that the 3’UTR 9’ emerged to confer on *tpm1* mRNA a translational control different from that conferred by the 3’UTR 9B. However, we found that the embryos injected with the pAMO had dramatic developmental defects. In control embryos, the neural folds elevate from the neural plate and progressively converge and fuse in a posterior to anterior direction to form the neural tube between stages 15 and 19 (Fig. 5B, left). In embryos injected with pAMO, the neural folds do not elevate or do not converge and fail to properly form the neural tube (Fig. 5B, right). To confirm that this phenotype results from a detrimental accumulation of 3’UTR 9B, we injected an mRNA encoding luciferase and containing the 3’UTR 9B. The injected embryos had the same developmental defect. Quantification of these phenotypes demonstrated that more than 90% of the embryos injected with the pAMO, and 80% of the embryos injected with luciferase-9B mRNA had defective neural folding. This was considerably more than embryos injected with a control morpholino, or embryos injected with a luciferase RNA containing the 9’ or a globin 3’UTR (Fig. 5C). At older stages, the tadpoles injected with the pAMO had a curved and bent body (Fig. S6). They were virtually unable to move, even when tickled with a needle. Their travel distances were as low as those of tadpoles injected with a *tpm1* translation-inhibiting morpholino that depletes them of muscular tropomyosin (Fig. 5D). Together, these data suggest that accumulation of high amounts of exon 9B-containing mRNA is detrimental at different developmental stages.

**Figure 5.**
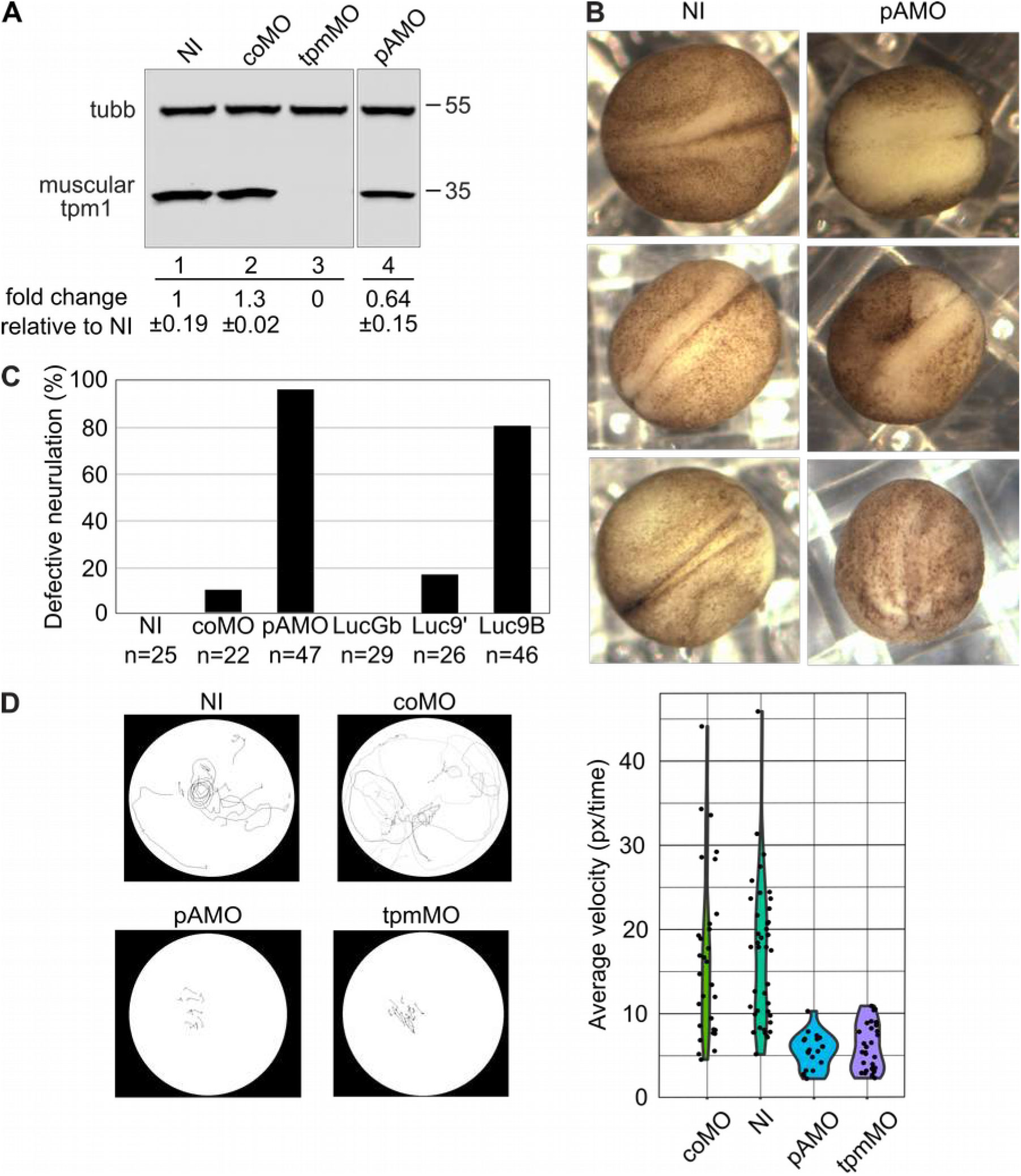
Phenotypes of embryos with high amounts of exon 9B. **A**, We injected two-cell embryos in both blastomeres with the indicated morpholinos and allowed them to develop until stage 28. We extracted proteins and analyzed them by western blot with antibodies against muscular tropomyosin or beta-tubulin as a loading control. tpmMO, translation-blocking morpholino against *tpm1*; (NI), non-injected. The quantification of 3 independent experiments is shown below the blot. Data is represented as mean ± standard deviation. **B**, We injected two-cell embryos in both blastomeres with the pAMO and allowed them to develop until stage 17. Representative photographs of neurulae are shown. The alterations in pAMO-injected embryos consisted of defective elevation of the neural folds or defective convergence and closure of the neural folds. **C**, quantification of defective neurulation following injection of the indicated molecules. **D**, Velocity of tadpoles injected with the indicated morpholinos. Left, movement tracking. We placed tadpoles in Petri dishes and recorded their movement during a fixed time period. Right, quantification of 30 movement tracking experiments. Velocity is expressed as pixels traveled by the tadpoles per unit time.

## Conclusion

Excessive accumulation of mRNA containing *tpm1* exon 9B is detrimental for *Xenopus laevis* development. It alters neurulation in early stages, and strongly reduces larval motility at later stages. Importantly, these defects are not only caused by the direct injection of an exon 9B-containing mRNA, but also by the injection of a morpholino that switches the endogenous *tpm1* maturation pattern from 8-9A9’ to 8-9A-9B. Hence, the observed developmental defects arise from the accumulation of exon 9B in somites that endogenously express isoform 8-9A9’. How does accumulation of exon 9B in somites interfere with neurulation? Neural tube patterning is achieved by crosstalk between four signaling pathways: FGF, Hedgehog, retinoic acid and BMP, originating from tissues in close proximity to the neural plate (Wilson and Maden 2005). Some of these signaling molecules originate from the somites. It is tempting to speculate that exon 9B acts as a sponge to titrate regulatory molecules present in somites, presumably RNA-binding proteins and/or microRNAs, resulting in altered signaling from the somites. A full description of the alterations to signaling pathways in *Xenopus* larvae with high amounts of exon 9B and the corresponding consequences for neurulation will require further experiments.

It appears that the toxicity of exon 9B in somites was at least part of the evolutionary pressure that led to the emergence of an alternative 3’UTR for muscular *TPM1* in sarcopterygians. The genomic region immediately downstream of a previously strictly internal exon underwent exonization resulting in a compound internal (9A) and terminal (9A9’) exon. The definition of terminal exon 9A9’ relies on the UTE and on a weak cleavage-polyadenylation site. RNase protection assays demonstrated that the UTE assists recognition of the weak cleavage-polyadenylation site, thereby coupling splicing with cleavage and polyadenylation. Interactions of splicing factors with cleavage and polyadenylation factors have been described, for example between U2AF65 and the carboxy terminal region of the poly(A) polymerase (Vagner et al. 2000), cleavage and polyadenylation specificity factor (CPSF) (Kyburz et al. 2006), or cleavage factor I CF1(m) (Millevoi et al. 2006). The UTE is in close vicinity to the branch point, which binds the U2 snRNP. This suggests that as yet unknown UTE-binding factors can interact with the U2 snRNP and cleavage-polyadenylation factors to improve the crosstalk between the 5’ and 3’ boundaries of exon 9A9’. This raises the question: how widespread is exonization caused by the appearance of a functional cleavage-polyadenylation site in an intron? Intron 8 has two unusual properties: it is AG-independent, and the branch point is far from the 3’ splice site. This large region between the branch point and the 3’ splice site had a great potential for the appearance of new intronic cis-regulators of pre-mRNA maturation, and the UTE is precisely in that region. Accordingly, exonization by the emergence of an active cleavage-polyadenylation site might be favored downstream of AG-independent introns with a remote branch point. It will be important to systematically assess whether exonization mediated by the appearance of an active cleavage-polyadenylation site in an intron is frequent close to introns with similar properties.

## MATERIALS AND METHODS

### Comparative genomic analysis

Sequence data for *TPM1* ortholog genes were collected for 30 species representing ray-finned fishes, sarcopterygians and amniotes. All the sequences, annotations and accession numbers are given in Table S1. When available, synteny was evaluated using the *TLN2* and *LACTB* genes surrounding the *TPM1* locus. Exonic annotation was manually curated for each species and is reported in Table S1. The properties of the branch points, 5’ splice sites, and cleavage-polyadenylation signals were evaluated using BPfinder (Corvelo et al. 2010), MaxEntScan (Yeo and Burge 2004), and polyA_SVM (Cheng et al. 2006). UTE scoring was performed using PWMEnrich in R (Stojnic and Diez 2021).

### Plasmids construction

The pBS-Actin-tpm1a-7-9B splicing reporter gene has been described previously, along with reporter genes with mutant 5’ or 3’ splice sites (Hamon et al. 2004). The transcription of the *X. laevis tpm1* genomic region from exon 7 to exon 9B was driven in embryonic myotomal cells by the cardiac actin promoter. In compound reporter genes, the region between the SnaB1 (20 nucleotides downstream of exon 8) and Kpn1 (310 nucleotides downstream of exon 9A) sites was replaced by equivalent sequences from other species, between around 600 nucleotides upstream and around 300 nucleotides downstream of exon 9A. For luciferase reporter plasmid construction, we inserted exon 9A or 9’ downstream of the firefly luciferase gene in the pmirGlo vector (Promega).

### *Xenopus laevis* embryo manipulations

*X. laevis* embryos were obtained by *in vitro* fertilization of eggs from laboratory-reared females. Embryo injections were carried out as described previously (Hamon et al. 2004). For splicing reporter gene injection, 250 pg of supercoiled DNA were injected into one blastomere at the two-cell stage. The sequences of the morpholinos are given in Table S2. We injected 0.5 pmol (bpMO) or 2 pmol (other morpholinos) into both blastomeres of two-cell embryos.

Luciferase RNAs were obtained by *in vitro* transcription from SalI-linearized vector using the mMessage mMachine T7 kit (Ambion). We injected 1 fmol of Luciferase mRNA into one blastomere of two-cell embryos and the embryos were cultured until Nieuwkoop and Faber stage 17 or 30. Proteins were extracted as previously described (Le Sommer et al. 2005), diluted in a luciferase assay buffer (PBS, 0.1% BSA) and mixed with furimazine (0.25% Nano-Glo® Substrate, Promega). The samples were distributed in duplicates in 96 half-well white polystyrene microtiter plates. Luminescence was measured using a Xenius XL luminometer (SAFAS).

For embryo tracking, pools of about 12 stage 40 embryos injected with the indicated Morpholinos were collected for tracking analysis. Upon settlement of the embryos, displacement was initiated by addition of one drop of 50 % ethanol to the water. Movies were recorded on a canon EOS 1100D for 10 seconds as “. Mov” files. Individual png images were recovered every 0,1 seconds from the video file using FFmpeg (Tomar 2006). Image were imported as 8-bits gray scale image stack to Fiji and individual embryos were tracked using the manual tracking plugin to record displacement over time (Schindelin et al. 2012). For each embryo, total displacement was normalized to recording time to compute a pixel/time average speed.

### Isolation of RNA, analysis by RT-PCR, RT-qPCR and RNase Protection Assay

RNA extracted from embryos was analyzed by an RT-PCR assay as described previously (Anquetil et al. 2009). PCR amplification was performed with a ^32^P-labelled forward primer in *tpm1* exon 8 (endogenous transcript), or a ^32^P-labeled forward primer at the actin promoter/exon 7 junction (splice reporter gene), and reverse primers in exons 8, 9’ or 9B. Amplimeres were resolved on 8% polyacrylamide gels and quantified by PhosphorImager analysis (Amersham Biosciences). Real-time PCR was performed using the Power SYBR green PCR master mix in an ABI Prism 7900 (Applied Biosystems). The RNA probes used for the detection of endogenous *tpm1, eef1a1* and *myl1* transcripts, and cleaved and uncleaved *tpm1* transcripts were described previously (Anquetil et al. 2009). RNase protection assays were performed using 10 µg of total RNA as described previously (Le Sommer et al. 2005). The sequences of the oligonucleotides used for RT-PCR and RT-qPCR are given in Table S2.

### Antibodies and Western Blot Analysis

Western blot analysis was carried out as described previously (Anquetil et al. 2009). Blots were detected with the Odyssey infrared imaging system (Li-COR Biosciences) and the following primary antibodies: monoclonal anti-tropomyosin clone TM311 (catalog number T2780, Sigma) and monoclonal anti-beta tubulin clone AA2 (catalog number T4026, Sigma).

## ACKNOWLEDGEMENTS

The authors gratefully acknowledge the help of Stéphane Deschamps, Margaux Véron and Claire Bertrand with the experiments. This work was supported by a grant from AFM-Telethon to SH.

## DATA AVAILABILITY

The data underlying this article are available in the article and in its online supplementary material.

