## Supplemental figures for "EXONIZATION BY THE EMERGENCE OF A CLEAVAGE-POLYADENYLATION SITE"

### SUPPLEMENTARY MATERIALS

#### Legends to supplementary tables

**Table S1. Genomic references**

Ensembl reference IDs of the *TPM1* genomic loci for the indicated species. The start, end and genomic strand of the indicated exons are given.

**Table S2. Oligonucleotide and morpholino sequences**

Sequences, description and usage of the oligonucleotides and Morpholino oligonucleotides used througout the study.

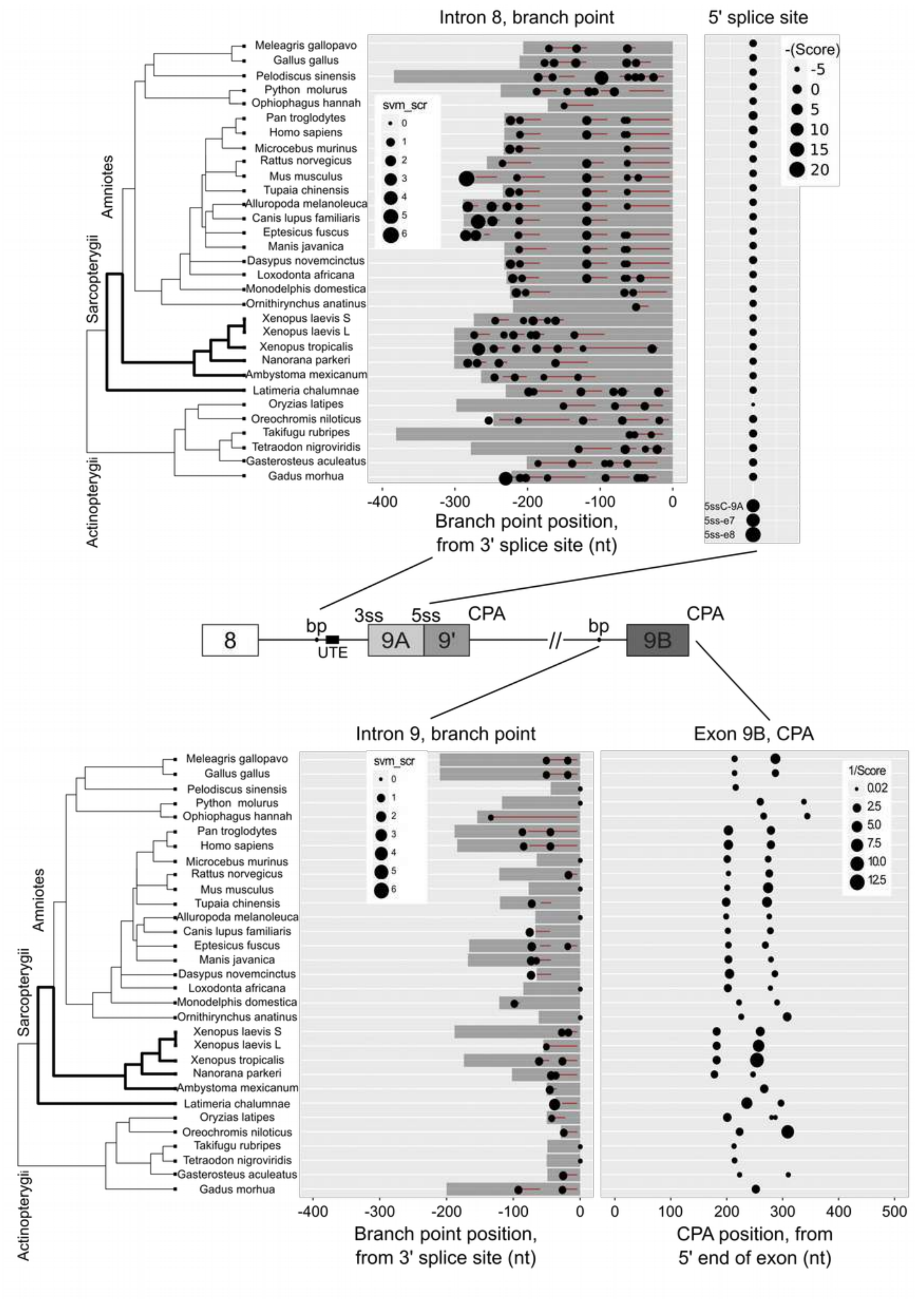


**Figure S1. Scoring cis-sequences potentially involved in *TPM1* pre-mRNA maturation**

Same analysis as in Fig. 1C, but the following cis-sequences were scored: branch point in intron 8, 5' splice site between exonic regions 9A and 9', branch point in intron 9, and cleavage-polyadenylation site in exon 9B. For the branch points, the red lines indicate the polypyrimidine tracts and the dark regions are the AG exclusion zones. The scores of other 5' splice sites in *Xenopus laevis* are shown for reference (5ssC-9A, consensus 5' splice site downstream of exon 9A as in Fig. 2B lanes 4, 11; 5ss-e7 and 5ss-e8, 5' splice sites downstream of exons 7 and 8). They are stronger than the weak 5' splice site downstream of exon 9A.

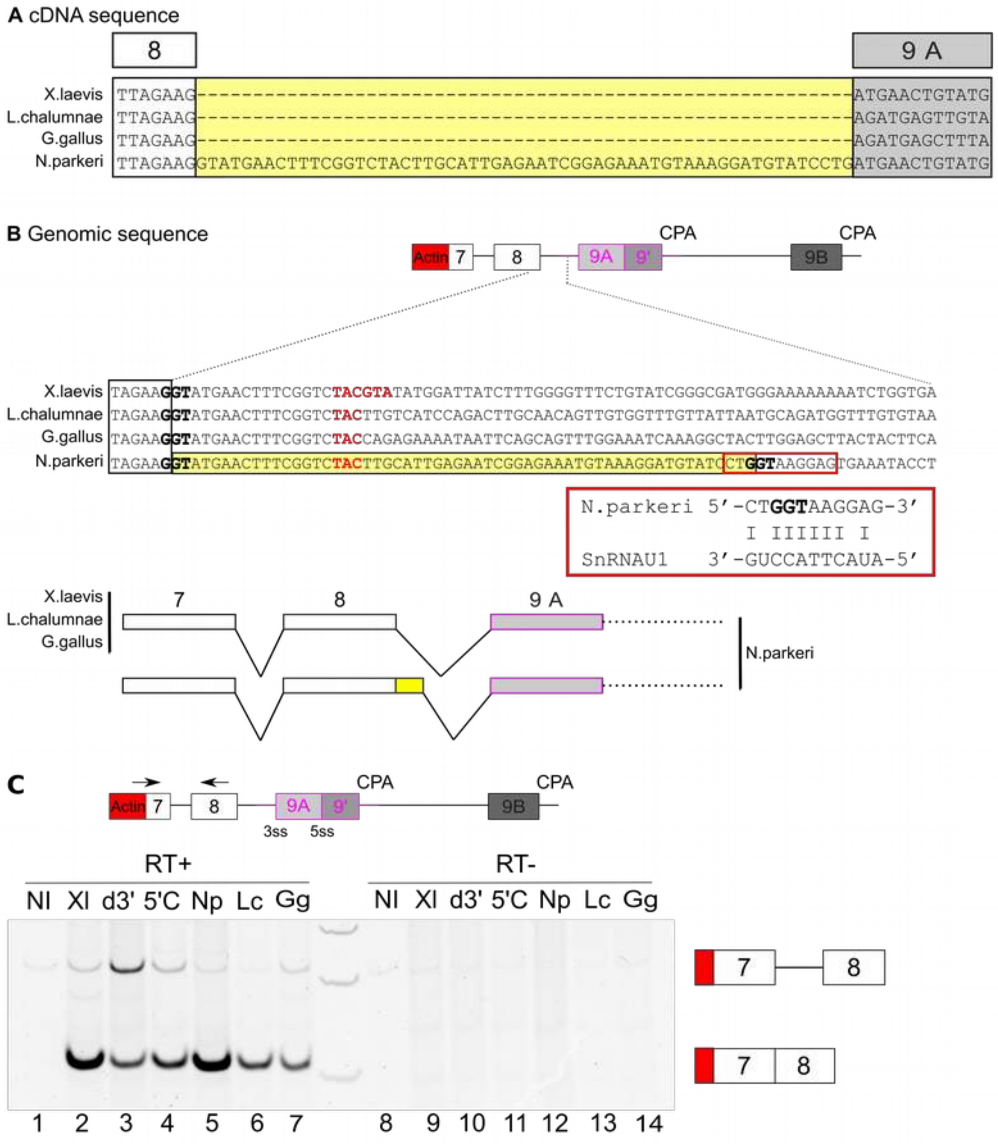


**Figure S2. A cryptic 5' splice site in *Nanorana parkeri* intron 8**

We excised the bands from the gels shown in Fig. 2B and we sequenced them. **A**, Alignment of the sequences of the bands obtained from the *X. laevis* splicing reporter gene and from 3 different gene compound reporter genes (*X. laevis* and the indicated species, bands indicated by a star in Fig. 2B for the *X. laevis* - *N. parkeri* compound gene). All the sequences align with exons 8 and 9A, except the sequences obtained from the *X. laevis* - *N. parkeri* compound gene which includes an additional sequence, boxed yellow. **B**, Sequences of the splicing reporter genes. Upper panel, *X. laevis* reporter gene, three lower panels compound genes (*X. laevis* and the indicated species). The SnaB1 site in the *X. laevis* sequence that was used for cloning is in red. The additional sequence in the transcript from the *X. laevis* - *N parkeri* compound gene (yellow in A) corresponds to the 58 first nucleotides of the intron. This 58 nucleotide region ends with a potential binding site for the snRNA U1. We conclude that a cryptic 5' splice site was used for mRNA production. **C**, Analysis of exon 7 - exon 8 splicing in the splicing reporter assay. Same analysis as in Fig. 2B, but we used primers in exons 7 (forward) and 8 (reverse) (lanes 1-7). We also show the negative controls carried out in the absence of reverse-transriptase (lanes 8-14).

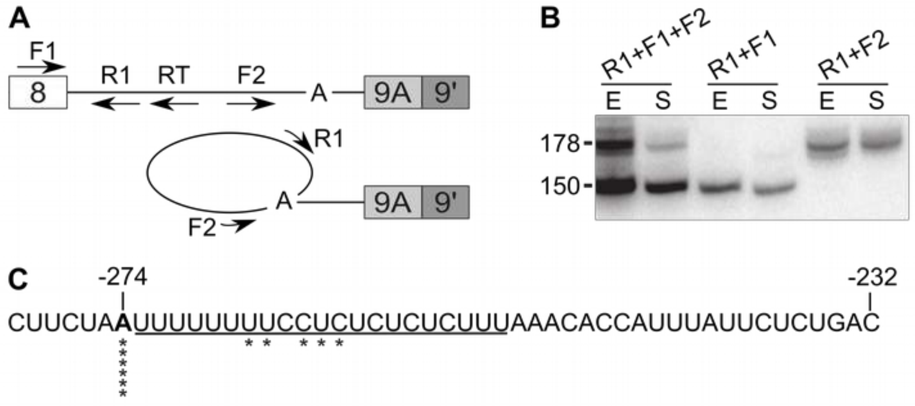


**Figure S3. Mapping the in vivo branch point on intron 8**

**A**, Positions of the primers used to identify intron 8 branch point site. RT is the primer for reverse transcription. R1, F1 and F2 are PCR primers. R1 and F1 amplify the cDNA obtained after reverse-transcription of *tpm1* pre-mRNA (150 nucleotides). R1 and F2 are divergent and can yield no amplimere from *tpm1* pre-mRNA, but they are convergent on intron 8 lariat and can yield an amplimere encompassing the branch point. Sequencing the amplimere identifies the branch point at a nucleotide-level resolution. **B**, RT-PCR products using RNA extracted from stage 28 whole embryos (E) and dissected somites (S), with the indicated primers. **C**, sequence of *tpm1* intron 8. The stars indicate the positions where the 5' splice site was ligated in the sequenced R1-F2 amplimeres. The underlined nucleotides are the polypyrimidine track. The A at position -274 is the branch point, while other nucleotides apparently ligated to the 5' splice site are very probably artifactual, arising from the RT crossing the ligated branch point.

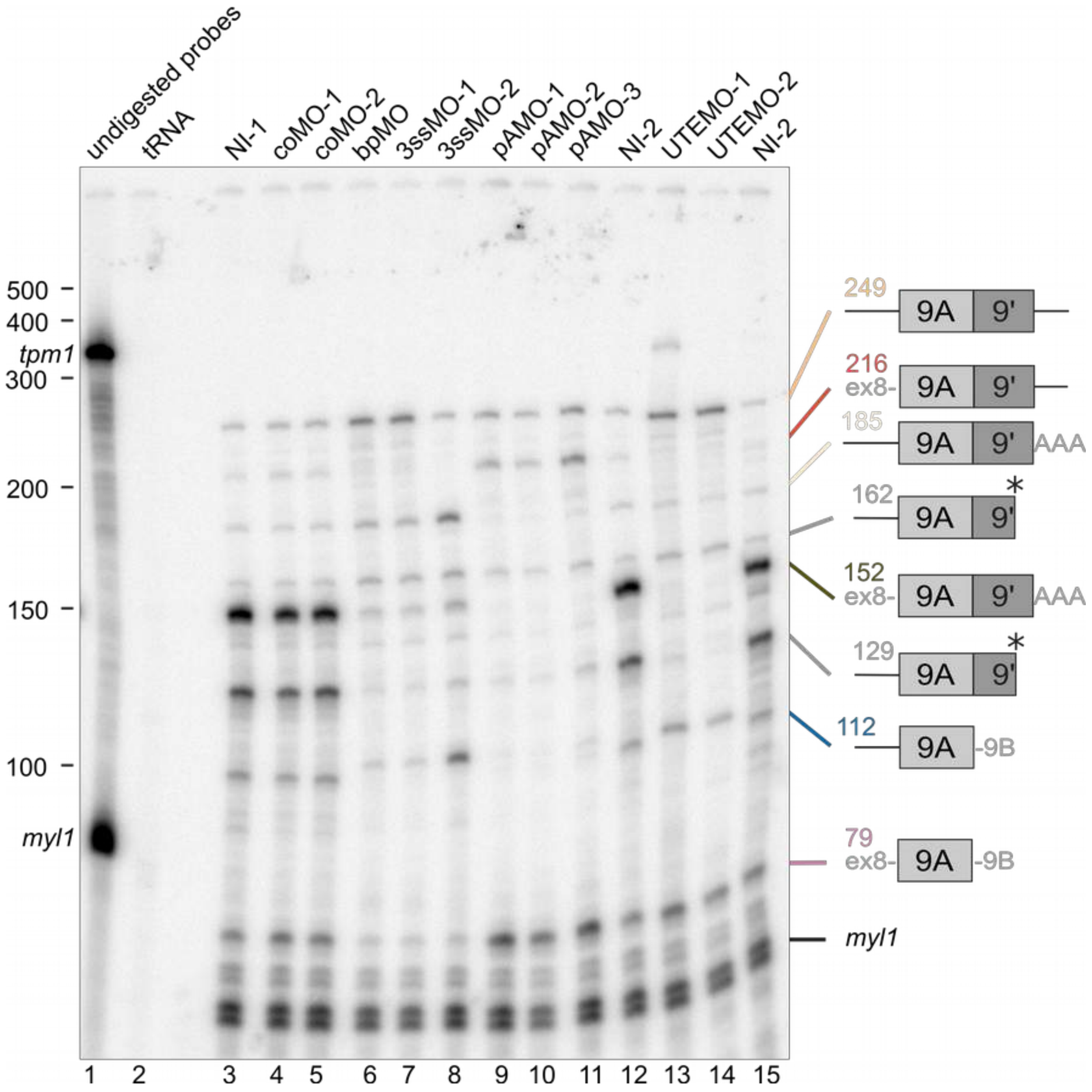


**Figure S4. Additional RNase protection assays experiments.**

Independent replicates of the experiment shown in Fig. 4.

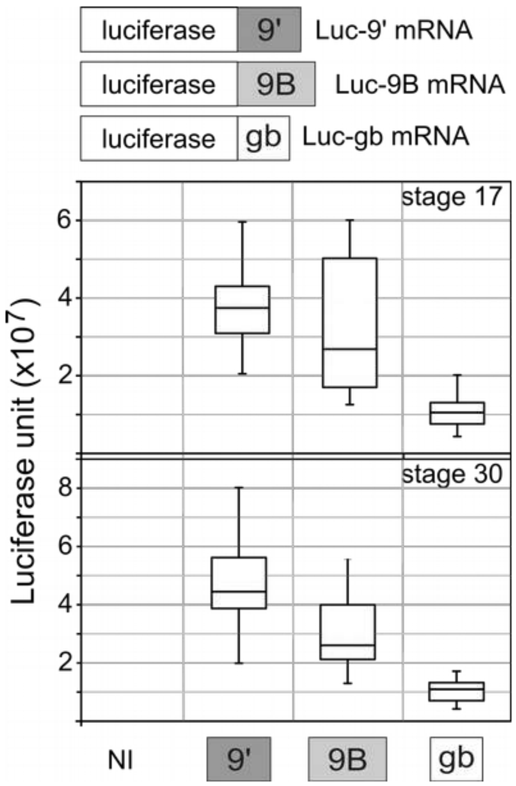


**Figure S5. Luciferase reporter assays.**

We injected two-cell embryos in both blastomeres with 1 fmol of luciferase mRNA containing the 9', 9B or globin 3'UTR, or we left them non-injected (NI). We allowed the embryos to develop until stage 17 or 30 as indicated We extracted proteins and measured luciferase activity. Twenty embryos were individually analysed in each condition.

\

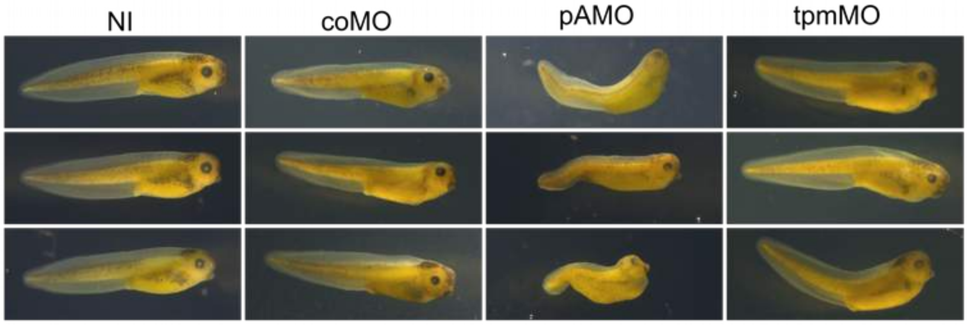


**Figure S6. Morphology of tadpoles with high amounts of endogenous exon 9B.**

Embryos were injected with the same morpholinos as in Fig. 5 and were allowed to develop until tadpole stage. We show here three representative photographs of the tadpoles for each condition.
